# A broad, exposome-type evaluation of xenobiotic phase II biotransformation in human biofluids by LC-MS/MS

**DOI:** 10.1101/2022.07.15.500202

**Authors:** Yasmin Fareed, Dominik Braun, Mira Flasch, Daniel Globisch, Benedikt Warth

## Abstract

Xenobiotics are chemicals foreign to a specific organism that humans are exposed to on a daily basis through their food, drugs and the environment. These molecules are frequently metabolized to increase polarity and subsequent excretion. During sample preparation, deconjugation of phase II metabolites is a critical step to capture the total exposure to chemicals in liquid chromatography mass spectrometry assays (LC-MS). Knowledge on deconjugation efficiencies of different enzymes and the extend of conjugation in human biofluids has primarily been investigated for single compounds or individual chemical classes. In this study, the performance of three β-glucuronidase and arylsulfatase mixtures from *H. pomatia*, from recombinant sources (BGS™), and from *Escherichia coli* combined with recombinant arylsulfatase (ASPC™) was compared and the efficiency of phase II deconjugation was monitored in breast milk, urine and plasma. An innovative LC-MS/MS biomonitoring method encompassing more than 80 highly diverse xenobiotics (*e.g*., plasticizers, industrial chemicals, mycotoxins, phytoestrogens, pesticides) was utilized for the comprehensive investigation of phase II conjugation in experiments investigating levels in breast milk and urine obtained from breastfeeding women. Overall, it was confirmed that *H. pomatia* is the most efficient enzyme in hydrolyzing different classes of xenobiotics for future exposome-scale biomonitoring studies. The recombinant BGS™ formulation, however, provided better results for breast milk samples, primarily due to lower background contamination, a major issue when employing the typically applied crude *H. pomatia* extracts. A deeper understanding of the global xenobiotic conjugation patterns will be essential for capturing environmental and food-related exposures within the exposome framework more comprehensively.

**Graphical abstract:** 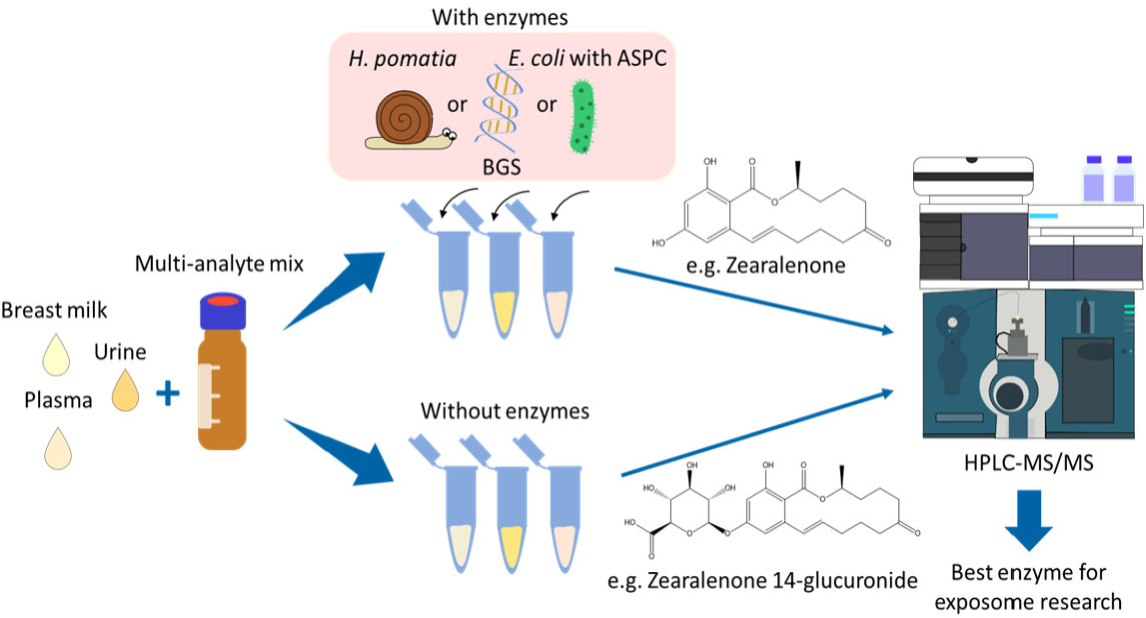

## Introduction

Humans are constantly exposed to thousands of xenobiotics on a daily basis. Xenobiotics are foreign compounds to the body that have been ingested and absorbed by individuals through external sources such as food, beverages, cosmetics, pollutants, and medications [1, 2]. Cumulatively referred to as the ‘exposome’, these exposures may compromise human health. The exposome describes the total amount of environmental exposures that the human body is subjected to in a lifetime [3, 4]. Typically, exposure is quantitatively assessed in biological samples (*e*.*g*., urine, blood, hair) by targeted or non-targeted human biomonitoring (HBM) approaches [3, 5]. Human biomonitoring is a rapidly growing field due to a rising concern for the chemical safety of people. In Europe, the European Human Biomonitoring Initiative (HBM4EU) was established in 2017 to increase public awareness regarding exposure to environmental and other anthropogenic chemicals [6]. A key tool to identify these environmental contaminants is liquid chromatography coupled to a tandem mass spectrometer (LC-MS/MS), where numerous metabolites can be simultaneously quantitated. It is also a highly sensitive platform that enables the detection of contaminants even below the ng/mL level [7].

Xenobiotics are metabolized and potentially activated by the human body and/or the microbiome. These contaminants are naturally lipophilic and are generally biotransformed into more polar, reactive compounds during phase I. In phase II, these compounds can be further conjugated with sugars, sulfates, glucuronides, and/or glutathione groups to increase the molecules’ hydrophilicity and facilitate excretion [2, 8]. Such biotransformation reactions are mediated by UDG-glucuronosyltransferases, N-acetyltransferases, glutathione S-transferases, methyltransferases, and sulfotransferases that are typically produced in the liver [9]. Potential biomarkers of exposure that can be analyzed in a given biological matrix frequently include conjugated derivatives such as glucuronides and sulfates [2, 10].

Phase II metabolites are commonly excreted from the body through urine; therefore most xenobiotics are found in its metabolized form in this matrix. Hence, urine is a crucial human sample for human biomonitoring, as the majority of human exposures can be determined in it [1, 8, 11]. Conjugated compounds are not restricted to urine and several studies reported phase II metabolites in human plasma [12–15]. Only a few studies suggest that there are also phase II conjugates in breast milk [16, 17].

The conjugated compounds can be analyzed directly by measuring the metabolized compound or following an enzymatic treatment step where the conjugates are cleaved and reverted to the native form. Reference standards of the biotransformed molecules are often not commercially available, thus hampering a direct approach for many xenobiotics of interest. As appropriate conjugated standards are not always available, an initial enzymatic hydrolysis step is usually utilized to quantitate total xenobiotics. This method is also more efficient as both non-conjugated and conjugated forms of the analyte can be simultaneously identified [12, 18, 19]. Most research into the detection of phase II metabolites utilizes a β-glucuronidase and sulfatase mixture from *Helix pomatia*. These enzymes are derived from the gastric juices of the Burgundy snail, but as a consequence of the biological nature and the subsequent presence of the inherent background matrix, these enzymes have posed challenges in research. In addition, the use of *H. pomatia* in xenobiotic hydrolysis has been questioned as being unethical [12]. Other enzymes such as β-glucuronidase from *Escherichia coli* and arylsulfatase from *Aerobacter aerogenes* have also been used for hydrolysis [12, 20, 21]. The snail *H. pomatia* produces not only β-glucuronidase, but also arylsulfatase, thus simplifying the simultaneous detection of both glucuronides and sulfates [12]. Another enzyme that is widely used in the detection of phase II metabolites is the β-glucuronidase from *E. coli*. According to previous studies, it appears to efficiently deconjugate isoflavones but has been criticized for the lack of arylsulfatase [12, 22]. A new alternative to potentially aid the simultaneous detection of glucuronides and sulfates is recombinantly produced enzymes. BGS™ recombinant is a recombinant enzyme of β-glucuronidase that also includes arylsulfatase. Additionally, a recombinant arylsulfatase (ASPC™ recombinant) was combined with *E. coli* β-glucuronidase. A recent study by Jain *et al*. compared the arylsulfatase from ASPC recombinant to the commercially available arylsulfatase from *H. pomatia*. Both enzymes detected similar quantities of metabolites, thus confirming the efficiency of the ASPC recombinant sulfatase [23].

The aim of the current study was to compare deconjugation efficiencies of mixtures containing β-glucuronidase and arylsulfatase, (i) from *H. pomatia*, (ii) recombinantly produced BGS and (iii) from *E. coli* with recombinant arylsulfatase (ASPC) with an LC-MS/MS method quantitating a multitude of xenobiotics and their respective conjugated forms in several biofluids. The most suitable enzymatic treatment for future rapid and straightforward deconjugation of each biofluid was then applied to a larger set of biological samples (urine, breast milk and plasma) and screened for 80+ xenobiotics prior to and after enzymatic hydrolysis to investigate the extend of conjugation in these biofluids.

## Materials and Methods

### Chemicals and Reagents

Assessed enzymes were *recombinant BGS* ^*TM*^ (β-glucuronidase and arylsulfatase mixture, Kura Biotech, Los Lagos, Chile), *E. coli* β-glucuronidase type IX-A (Sigma-Aldrich, Darmstadt, Germany), ASPC^*TM*^ recombinant arylsulfatase (Kura Biotech, Los Lagos, Chile), and *H. pomatia* β-glucuronidase/arylsulfatase mixture (Roche Diagnostics, Vienna, Austria). The substrate phenolphthalein β-D-glucuronide, sodium chloride, ammonium acetate, potassium phosphate, acetic acid, and citrate plasma were purchased from Sigma-Aldrich (Darmstadt, Germany) and 4-nitrocatechol sulfate from ChemCruz (Santa Cruz Biotechnology). Phenolphthalein standard solution, dipotassium hydrogen sulfate, sodium chloride, and glycine were purchased from Carl Roth (Karlsruhe, Germany). Bovine serum albumin (BSA), magnesium sulfate, and sodium hydroxide pellets were purchased from Fisher Scientific (Vienna, Austria). Instant buffer II was provided by Kura Biotech (Los Lagos, Chile). Ammonium carbonate and LC-MS grade acetonitrile were purchased from Honeywell (Seelze, Germany). LC-MS grade water was obtained from VWR chemicals (Vienna, Austria).

For determining the efficiency of the enzymes, the following reference glucuronide and sulfate conjugates were obtained from Toronto Research Chemicals (Toronto, Canada): quercetin-7-O-β-D-glucuronide, genistein-7-β-D-glucuronide, caffeic acid-3-β-D-glucuronide, daidzein-7-β-D-glucuronide, and genistein-7-sulfate. Kaempferol-3-O-glucuronide was obtained from Extrasynthese (Genay, France). Estradiol-17-glucuronide, p-cresol-sulfate, and estradiol-3-sulfate were purchased from Sigma-Aldrich (Vienna, Austria). The glucuronides and sulfates were dissolved in acetonitrile to a final concentration of 0.03–12 ng/mL and combined and stored at -20°C. For the exact concentration of each glucuronide and sulfate in the multi-analyte mix, refer to supplementary Table S3.

Reference standards of the 86 assessed xenobiotics were used for the quantitative assessment and are listed in the supplementary information. For detailed information such as MS/MS parameters, transitions, retention times, reference standard concentrations, as well as purchasing information please refer to Jamnik *et al*. (2022) [24]. The reference standards were combined and dissolved in 10% ACN at final concentrations of 0.3–100 ng/mL for subsequent use as standard calibrants. Due to availability, an internal standard solution with ^13^C_18_ Zearalenone (ZEN) (Cambridge Isotope Laboratories, Tewksbury, Massechusetts) was used for quality control purposes. The solution was prepared by adding 1 mL of ^13^C_18_ ZEN to 19 mL ACN:MeOH (1:1). All prepared reference standards and solutions were stored at -20°C until required [24].

### β-Glucuronidase Activity Assay

The three enzymes of interest with glucuronidase activity, recombinant BGS, *E. coli* glucuronidase without additional ASPC, and *H. pomatia* glucuronidase/arylsulfatase were tested on their β-glucuronidase activity for standardization of the enzyme activity in follow-up experiments. The procedure was conducted according to a vendor protocol [25]. For quantitation, phenolphthalein glucuronide calibration standards were prepared at a concentration range between 1 and 5 µg/mL and measured without prior heat treatment. Subsequently, all three enzymes were diluted to approximately the same activity level (700 U/mL) based on their stated individual activity in the certificate of analysis provided by the manufacturers. To determine the activities of the enzymes, 130 µL HPLC-grade water was added to 100 µL 75 mM potassium phosphate buffer. A volume of 50 µL of a defined phenolphthalein-glucuronide standard solution (0.05% w/v) was added and mixed in a micro reaction tube. This mixture was then equilibrated to 37°C (*H. pomatia)* or 45°C (*E. coli* and recombinant BGS) in a thermoshaker. Once the tubes reached the appropriate temperature, 20 µL enzyme solutions (∼700 U/mL) were added to the corresponding tubes and the mixtures were kept at 37°C (*H. pomatia*) and 45°C (*E. coli* and recombinant BGS) for exactly 30 minutes. Finally, 1 mL of 200 mM glycine buffer was immediately added to the samples to stop the reaction. As a negative control, the different enzymes were diluted 1:100 (v:v), treated in the same way, and included in the measurement. After transferring all samples, controls and standards to a 96-well plate, the solutions were measured at 540 nm in a plate reader. The phenolphthalein concentration was plotted against its absorbance at 540 nm. The final activities of the enzymes (in fishman units) were calculated using the following equation:

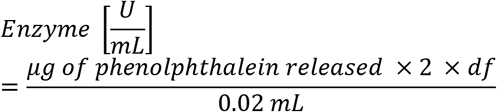

Where

2 = time correction of assay (Unit Definition is 1h)

df = dilution factor

0.02 = volume of enzyme used [mL]

#### Equation 1

Equation for the determination of the glucuronidase activities in U/mL after calculating the amount of released phenolphthalein from the measured absorbances

### Arylsulfatase Activity Assay

The sulfatase activity of recombinant BGS™, ASPC™ arylsulfatase, and sulfatase from *H. pomatia* was tested using a vendor protocol from Sigma Aldrich [26]. Firstly, 50 µL of the appropriate buffer for the specific enzyme (instant buffer II for recombinant BGS and ASPC arylsulfatase and 2.5 M acetate buffer for *H. pomatia*) was added to 40 µL of 6.25 mM p-nitrocatechol-sulfate solution. The solution was gently mixed and equilibrated to 37°C for the experiments using *H. pomatia* and 45°C for recombinant BGS and ASPC arylsulfatase, respectively. Next, 0 µL (process blank), 5 µL, 7 µL and 10 µL of each of the undiluted, 10-fold diluted, and 50-fold diluted enzyme were added for standard addition. These solutions were gently mixed and incubated at 37°C and 45°C accordingly for exactly 30 minutes. 500 µL of a 1 M sodium hydroxide solution was then immediately added to stop the reaction. The difference in added enzyme volumes was adjusted with the corresponding enzyme diutions to give a total of 10 µL enzyme solution in each sample. This was necessary to eliminate differences in signal intensities originating from the enzyme. After all solutions were transferred to a 96-well plate, the samples were measured at 515 nm in a plate reader. The activities of the enzymes in units/mL were calculated with the following equation:

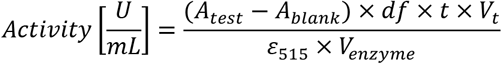

Where

A_test_ = Absorbance of the test solution at 515 nm.

A_blank_ = Absorbance of the blank solution at 515 nm.

df = dilution factor of the enzyme.

t = time correction factor of one hour.

V_t_ = total volume of the assay [mL].

ε_515_ = Milimolar extinction coefficient of p-nitrocatechol at 515 nm.

V_enzyme_ = volume of enzyme added to the solution [mL].

#### Equation 2

Equation used to determine the arylsulfatase activities of the assessed enzymes in U/mL.

### Samples

Pooled biofluid samples for matrix-matched calibration were obtained as follows: lithium-heparin plasma was purchased from Innovative Research (Novi, Michigan). Pooled urine was provided by an Austrian female volunteer who avoided the consumption of food and use of cosmetics stored in plastic containers, and food high in phytoestrogens for two days prior to sample collection. The Semmelweis Women’s Clinic in Vienna, Austria kindly provided pooled breast milk samples. These pooled samples were used for matrix-matched standards and were stored at -20°C for urine and breast milk samples and -80°C for plasma samples, prior to analysis.

For the proof-of-principle studies, well-characterized breast milk and urine samples from a previous investigation were used (Braun *et al*. 2020). These had been collected as follows: in the long-term longitudinal study (#1), a lactating mother pumped breast milk at home into sample containers. Samples that were collected on the same day, and in case of low available volumes across two days, were mixed to create an aggregated sample. The provided breast milk amount varied depending on the needs of the infant. A total of 87 breast milk samples were collected. The collected breast milk was stored in the refrigerator at 4°C for a maximum of two days before storage at -20°C until analysis. The short-term longitudinal study (#2) was conducted with three mothers aged between 25 and 32 years who manually collected breast milk every morning for five consecutive days. The mothers also collected the first morning urine during these five days. All three mothers collected samples two to three months *post-partum*. The women did not have any specific diet and all had normal body mass indices. The study was approved by the University of Vienna Ethics Committee and the Ethics Committee of Lower Austria, where the sampling occurred under the authorization number #00157 [27]. As no plasma samples were available, the commercial pooled lithium-heparin plasma was measured in triplicate to investigate also this human biofluid.

### Sample preparation

#### Hydrolysis Efficiency of Enzymes

For ideal conditions for the tested enzymes, the pH stability after the addition of several buffers was tested. For details refer to the Supplementary Information. In a preliminary experiment, the efficiencies of the three β-glucuronidase/arylsulfatase enzyme mixtures of interest from (i) *H. pomatia*, (ii) BGS recombinant, and (iii) *E. coli* with ASPC recombinant were tested by spiking the three matrices (breast milk, urine, and plasma) with a mixture consisting of eleven selected glucuronide and sulfate conjugate reference standards (refered to as ‘conjugate reference mix’). The three enzyme preparations were compared to an untreated sample for each matrix. Each condition was measured in triplicate. For samples digested with *H. pomatia*, ammonium acetate buffer (2.5 M) was used to control the pH; whilst for BGS recombinant and *E. coli* with ASPC recombinant the instant buffer II from KURA biotech was used. The extraction procedure for the three matrices used for this experiment as well as the proof-of-principal experiments are described below.

The hydrolysis efficiencies in supplementary table S9 was calculated using the ratio of the average peak height from the parent compound of the conjugate after hydrolysis with the tested enzyme mixture and the total height of the undigested control with the enzyme treated sample. This ratio was then multiplied by 100 to allow a percentage comparison of the individual values.

### Calibrants

To prepare the matrix-matched calibrants, 200 µL non-spiked pooled samples of each matrix were prepared as mentioned above according to its respective matrix and reconstituted in 200 µL of the respective solvent standards containing all 89 analytes together with nine selected conjugated glucuronides and sulfates. However, the enzymatic treatment was omitted in the preparation of the matrix-matched standards to avoid the release of additional background contamination.

### Plasma and Urine

Plasma and urine samples treated with BGS recombinant or *E. coli* with ASPC arylsulfatase were prepared based on the method from Preindl *et al*. (2019), adding an enzymatic treatment step. Briefly, the pH of the matrix samples (50 µL) was adjusted with 50 µL instant buffer II for samples treated with BGS or 50 µL instant buffer II with 4000 U of E. coli β-glucuronidase for samples treated with ASPC. Then, 20 µL of a standard stock mixture consisting of selected glucuronides and sulfates were spiked into the buffered biofluid samples for the pre-experiment. After adding 50 µL of the undiluted enzyme solution (BGS or ASPC), samples were incubated at 45°C for 16 h at 400 rpm. For the detailed activities of each enzyme mixture refer to supplementary Table S4. After incubation, 200 µL of extraction solvent (ACN:MeOH, 1:1, v/v) and 2.5 µL of ^13^C_18_ ZEN internal standard were added. The samples were then vortexed, sonicated for 10 min on ice and chilled at -20°C for 2 h to precipitate the proteins. Subsequently, samples were centrifuged for 10 min (18,000 × *g*, 4 °C) and the supernatant was transferred to a new micro reaction tube. Samples were dried overnight at 4°C in a vacuum concentrator (Labconco) and reconstituted with 50 µL of 10% ACN. After vortexing and centrifugation (10 min, 18,000 × *g*, 4°C), the supernatant was transferred to an amber glass vial and injected into the LC-MS system [28].

The sample preparation with *H. pomatia* enzymes was performed with 100 µL matrix with 20 µL of the standard stock mixture as mentioned above. Then, 100 µL of an enzyme solution (4000 U for the β-glucuronidase) in 2.5 M ammonium acetate buffer was added and incubated for 16 h with 400 rpm at 37 °C. The remaining extraction was performed as described above with reconstitution in 100 µL to align the dilution.

### Breast Milk

Breast milk samples treated with BGS recombinant and *E. coli* with ASPC arylsulfatase were prepared according to Braun *et al*. (2018) with slight modifications [29]. Samples (50 µL) were spiked with 20 µL of the glucuronide/sulfate mixture (conjugate reference mix) for the pre-experiment and after vortexing, 50 µL of the enzyme solution together with 50 µL of instant buffer II was added and incubated at 45°C for 16 h in a thermoshaker (400 rpm). 50 µL ACN fortified with 1% formic acid and 2.5 µL of ^13^C_18_ ZEN internal standard was then added and the samples were vortexed for 3 min. For phase separation, 20 mg of anhydrous magnesium sulfate and 5 mg of sodium chloride was added, the samples were vortexed for 3 min, centrifuged (10 min, 18,000 × *g*, 4°C) and the upper layer was transferred to new micro reaction tubes. Subsequently, samples were chilled at -20°C for 2 h to precipitate the proteins. After centrifugation (10 min, 18,000 × *g*, 4°C), the remaining supernatant was transferred to new micro reaction tubes and dried overnight in a vacuum concentrator at 4°C. Samples were then reconstituted in 10% ACN (50 µL), centrifuged (10 min, 18,000 × *g*, 4°C), transferred into glass vials and injected into the LC-MS system.

In the case *H. pomatia* enzymes were used, the sample preparation protocol differed slightly. 100 µL samples were spiked with 20 µL of the glucuronide/sulfate mixture (conjugate reference mix) for the pre-experiment and vortexed. 100 µL of an enzyme solution in 2.5 M ammonium acetate buffer (4000 U for the β-glucuronidase) was added, incubated at 37°C for 16 h in a thermoshaker (400 rpm). The following steps were conducted as described above with reconstitution in 100 µL to keep the dilution factor.

### LC-MS/MS Parameters and Analysis

Measurements were conducted using a validated method from Jamnik *et al*. (2021) [24]. A 1290 Infinity II LC system from Agilent coupled to a QTrap 6500+ mass spectrometer with a Turbo-V™ ESI source from Sciex was applied, utilizing fast polarity switching in multiple reaction monitoring mode (MRM). For the determination of the hydrolysis efficiency the MS/MS parameters and transitions of additional glucuronides and sulfates were derived from Oesterle et al. [30] and added to the method and are described in supplementary Table S1. To attain chromatographic separation, an Acquity HSS T3 (Waters) reversed-phase column (1.8 μm, 2.1 mm × 100 mm) together with a VanGuard pre-column (1.8 µm) from Waters was used. 0.3 mM of ammonium fluoride in LC-MS grade water was used as eluent A and acetonitrile was used as eluent B. 5 µL of sample was injected into the system at a flow rate of 0.4 mL/min. The column compartment and autosampler were maintained at constant temperatures of 40°C and 7°C, respectively. Elution of the analytes was achieved with the following gradient: 5% B from 0-1 min, then rise to 18% B until 1.8 min, followed by a further increase to 35% B until 4.2 min, then ascend to 48% B which remained constant until 13 min, and a final increase to 90% B until 15.8 min. The column was subsequently washed with 98% B until 17.6 min and recalibrated with 5% B from 17.7 min to 20 min.

### Quantitation, Data Evaluation, and Software

Evaluation of the data was performed using the Multiquant software (v3.0.3, Sciex). Each linear regression curve was performed with matrix-matched calibration standards to consider possible matrix effects and retention time shifts in the various matrices. The matrix-matched standards were measured after every 20^th^ sample and were used as linear calibration curve with a weighting of 1/x. The results were corrected using the average extraction efficiency of each analyte according to Jamnik *et al*. (2020) [24]. To account for contamination that may occur during the sample preparation procedure from reaction tubes, gloves, pipette tips, or the LC system, the unknown samples were also corrected with process blanks. These blanks were prepared by conducting the sample preparation protocol for each matrix (as described above) with water and quantitating against the solvent calibration curve. Contamination of the matrix itself was also considered through corrections with a matrix blank. This was conducted by preparing the breast milk, urine, and plasma sample used for the matrix matched calibration according to the sample preparation protocol above without enzyme or temperature treatment. Further calculations and data evaluations were carried out using Microsoft Excel 2020. For data acquisition, Analyst 1.7.1 from Sciex was used.

OriginPro (version 2021, OriginLab) was used for statistical analysis and to plot graphs. Figures were created in Inkscape (version 1.0.1, Inkscape) and chemical structures were generated with ChemDraw (version 12.0.2, PerkinElmer).

## Results

### Glucuronidase and Sulfatase Activity Assays

To standardize the enzyme activity for the experiments, activity assays were conducted for optimal comparison between the three enzyme mixtures (Table S7). According to the β-glucuronidase activity assay, the enzyme with the highest glucuronidase activity was the β-glucuronidase from BGS recombinant followed by *E. coli* glucuronidase and finally *H. pomatia*. All three enzymes were able to deconjugate phenolphthalein-β-glucuronide solution sufficiently. The enzymes were diluted for further experiments with the appropriate buffers; 2.5 M acetate buffer for *H. pomatia* and instant buffer II for BGS recombinant and *E. coli* with ASPC to reach a total of 2000 U/mL in the sample according to the resulting activities from the β-glucuronidase activity assay.

For the sulfatase activity on the other hand, *H. pomatia* was able to deconjugate p-nitrocatechol-sulfate at a 50-fold dilution displaying a sufficient digestion activity of 656 U/mL. Several isoforms of glucuronidase and sulfatase are present in the crude *H. pomatia* enzyme mixture, which likely affects the hydrolysis of selected glucuronides and sulfates. As shown in supplementary Table S7, the arylsulfatase from the recombinant BGS and ASPC enzymes were initially barely able to deconjugate p-nitrocatechol with the two tested buffers PBS and ammonium carbonate buffer, displaying an activity of merely 0.75 U/mL for BGS recombinant and 0.88 U/mL for ASPC recombinant.. After switching the buffer to the instant buffer II at the same conditions as with the other two buffers, the enzymes were able to deconjugate p-nitrocatechol-sulfate. Nevertheless, the recombinant enzymes showed much lower activities than *H. pomatia*, only able to deconjugate p-nitrocatechol-sulfate at a 10-fold dilution, with a significantly lower activity compared to its glucuronidase activity. Hence, the sample preparation for these enzymes was altered to match their sulfatase activities in all samples for ideal comparison. The enzyme mixture *H. pomatia* and *E. coli* was kept matching the glucuronidase activity of 2000 U/mL β-glucuronidase with 24.5 U/mL arylsulfatase activity in *H. pomatia*, while the recombinant enzymes were added at the same amount as buffer and sample for optimal hydrolysis (*e*.*g*., 50 µL matrix with 50 µL instant buffer II and 50 µL enzyme), at activities of 6.03 U/mL arylsulfatase and 212703 U/mL β-glucuronidase for BGS and 7.8 U/mL arylsulfatase for ASPC in the samples (supplementary table S4). To have reasonably high arylsulfatase activity in the BGS experiment, the β-glucuronidase was not diluted to an equal activity but the highest possible concentration for arylsulfatase achieved. This activity only had a 4-fold difference in arylsulfatase activity in comparison to the *H. pomatia* enzyme mixture. This was done by adding the enzyme at a ratio of 1:1:1 (sample:enzyme:buffer).

### Enzymatic Hydrolysis

In the preliminary experiment where the three enzyme mixtures were assessed on their hydrolysis efficiencies, the digested samples of all three matrices (urine, breast milk, and plasma) were analyzed in triplicate to determine the most promising enzyme for deconjugating phase II metabolites. The resulting signals were evaluated and the signal decrease for the conjugated analytes as well as the increase in the deconjugated state was assessed (refer to Figure 1).

**Figure 1.**
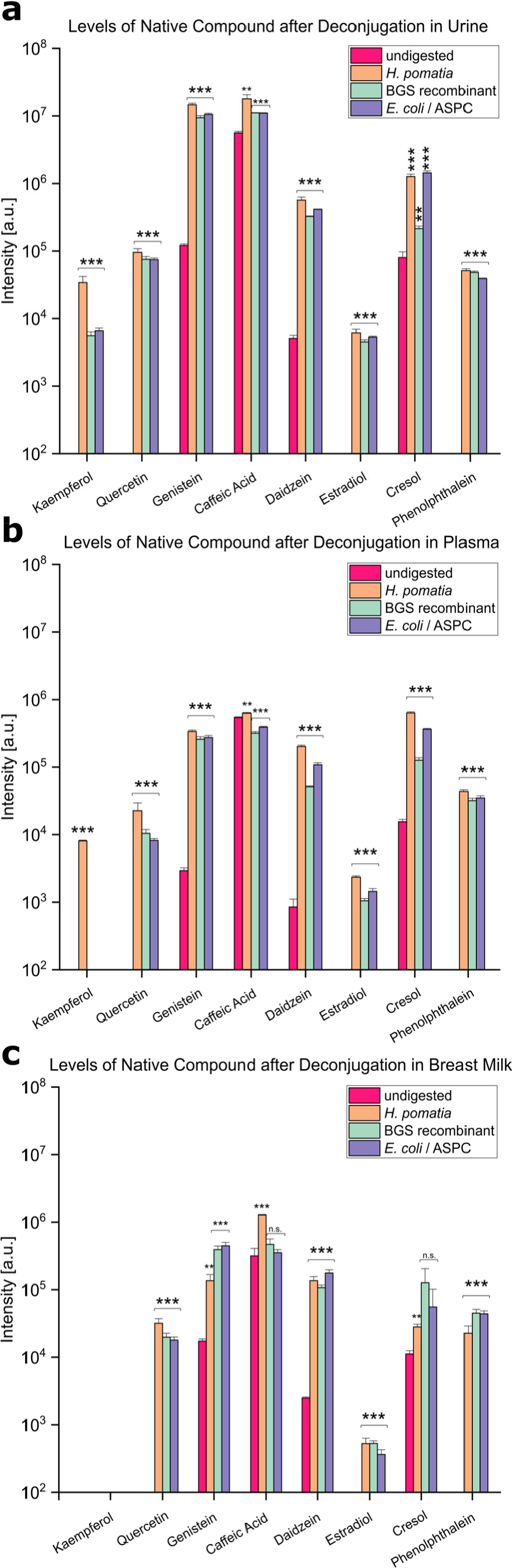
Level of deconjugated compound after hydrolysis for 16 h with the three enzyme mixtures assessed: *H. pomatia*, BGS recombinant, and a mixture of *E. coli* with ASPC recombinant for (a) urine, (b) plasma, and (c) breast milk. Statistical significance was shown with a student t-test against the undigested control, p values below 0.05 were considered significant (n.s. p > 0.05, * p < 0.05, ** p < 0.01, *** p < 0.001).

In all three matrices, it was evident that the free forms of kaempferol-3-O-glucuronide, quercetin-7-O-β-glucuronide, estradiol-17β-glucuronide, estradiol-3-sulfate, and phenolphthalein-β-D-glucuronide could only be detected after enzymatic hydrolysis. This highlights the importance of cleaving the glucuronide and sulfate conjugates when investigating xenobiotics in human samples. For all these eleven tested analytes, *H. pomatia* enzymes yielded the highest signals of deconjugated substances after hydrolysis (Figure 1). This is especially evident in the conducted student t-test comparing *H. pomatia* with the recombinant BGS and the mixture of *E. coli* with ASPC enzymes in Table S8. Here, all analytes significantly increased in plasma, except for quercetin with no significance. Genistein, daidzein, and cresol were all present at higher levels for all three enzyme preparations after deconjugation, despite already being present before enzymatic treatment. The student t-test conducted on these analytes were all significant in comparison with the undigested control, with the exception of cresol for the recombinant enzymes, BGS and *E. coli* with ASPC in breast milk, due to its high standard deviation, as illustrated in Figure 1. For plasma and urine samples, *H. pomatia* treatment resulted in the highest signals of free compounds, hydrolyzing eight analytes more sufficiently than the other two tested enzymes. However, when looking at breast milk, the two recombinant enzymes were able to deconjugate six analytes more rapidly than *H. pomatia*. Overall, despite varying enzyme activities, all three tested enzymes mixtures were able to hydrolyze most conjugates sufficiently in comparison to the undigested control, as evidenced by the significance illustrated in Figure 1. *H. pomatia* and BGS recombinant were selected for further experiments as the enzyme mixture ASPC with *E. coli* had no significant advantages (supplementary Table S8) in hydrolysis efficiencies compared to BGS, and required the combination of two enzymes.

### Background Contamination of Enzymes

In the experiments, signals for some of the 86 assessed xenobiotics were observed in the pure diluted enzyme solution. The contaminations in the respective enzymes used in further experiments, namely *H. pomatia* and BGS recombinant were quantified with the solvent standard calibration curve and illustrated in Figure 2. Contamination was found for 38 of the analyzed xenobiotics, with most of the contamination deriving from plasticizers, phytoestrogens, and personal care products. *H. pomatia* was observed to be contaminated with 37 xenobiotics, while BGS recombinant only contained 28 xenobiotics (supplementary Table S15).

**Figure 2.**
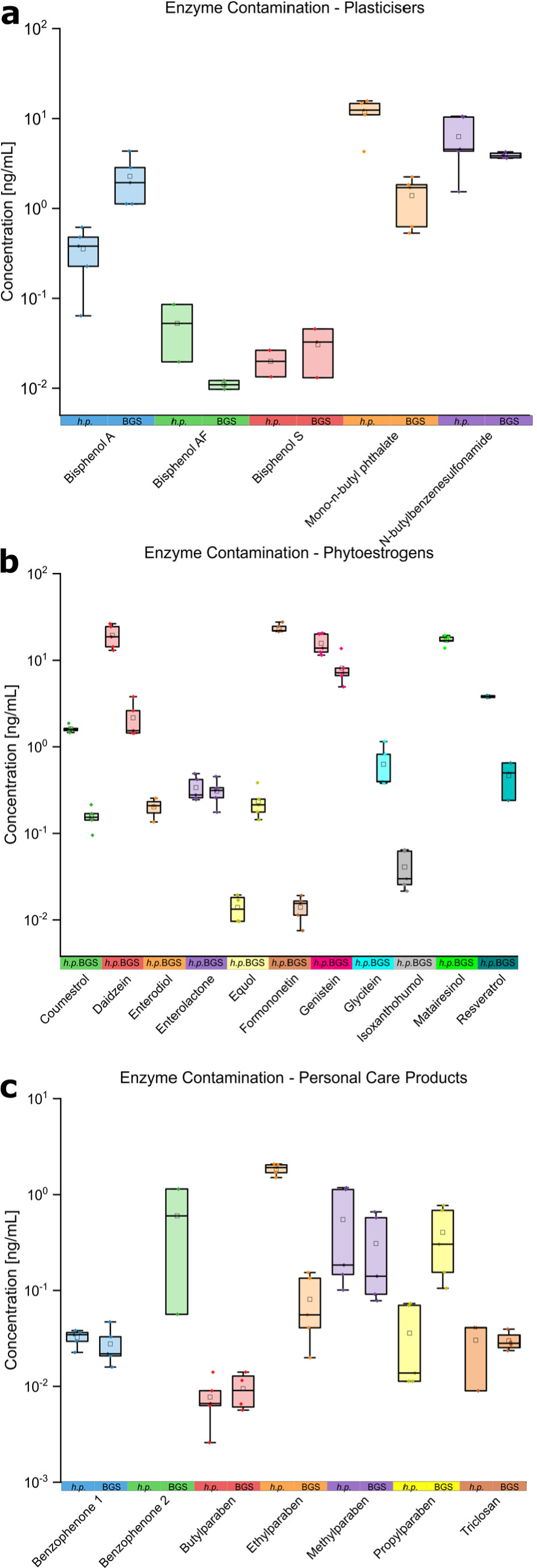
Background contamination signals observed in the two selected enzymes, *H. pomatia* extract (*h*.*p*.) and BGS recombinant (BGS) for (a) plasticizers, (b) phytoestrogens, and (c) personal care products. The black line in the box represents the median, the small square the mean, the box the 25–75 percentiles, and the whiskers the interquartile range. In the table, the minimum (min) and maximum (max) concentration in ng/mL are given together with the average (avg). To improve visibility the data was plotted on a logarithmic y-axis.

For most classes, the concentration of contamination in the enzyme mixture from *H. pomatia* was higher than in BGS recombinant. Only certain analytes, such as bisphenol A, bisphenol B, perfluorooctanoic acid, equol, glycitein, benzophenone 1, benzophenone 2, butylparaben, and propylparaben had slightly higher levels of contamination in BGS recombinant.

### Application to Urine

Urine samples were analyzed as part of the short-term breast milk study, where three different mothers collected their first morning urine on the same five days as the breast milk samples (n=15) (Figure 3). In total, 59 of the 85 xenobiotics were detected in the urine samples (Table S13), where 17 additional analytes were only present after enzymatic hydrolysis. In the urine samples, detected xenobiotic concentration levels significantly increased after enzymatic hydrolysis.

**Figure 3.**
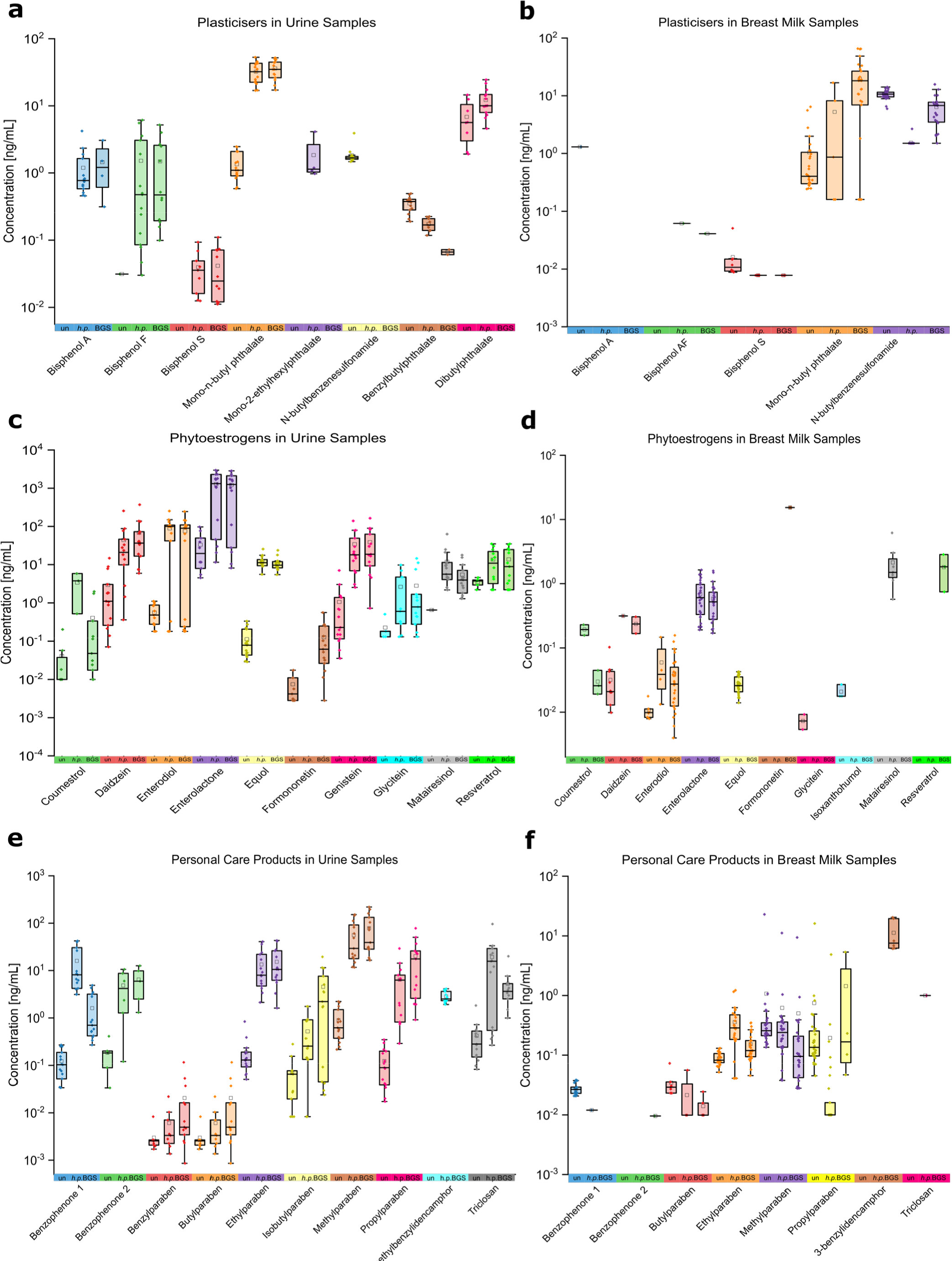
Results of the short-term longitudinal breast milk study, where three mothers collected breast milk (a, c, e) and urine samples (b, d, f) over five days (n=15) for (a, b) plasticizers, (c, d) phytoestrogens, and (e, f) personal care products. These samples were measured without enzymatic hydrolysis (un), and with an enzymatic treatment of *H. pomatia* (*h*.*p*.) at 2000 U β-glucuronidase and 24.5 U arylsulfatase and BGS recombinant (BGS) at 212703 U β-glucuronidase and 7.8 U arylsulfatase. The black line in the box represents the median line, the small square the mean, the box the 25–75 percentiles, and the whiskers the interquartile range. In the table, the minimum (min) and maximum (max) concentration in ng/mL are given together with the average (avg) and number of positive samples (n). The y-axis is displayed on a decadic logarithmic scale.

For plasticizers, *H. pomatia* was significantly more efficient than BGS recombinant in hydrolyzing the xenobiotic-conjugates, resulting in higher xenobiotic concentrations in six out of eight detected plasticizers. Despite the higher background contamination levels of *H. pomatia* in comparison to BGS recombinant, both enzymes mostly achieved similar hydrolysis efficiencies of the phytoestrogens and personal care products. The concentration of 28 out of 30 plasticizers, phytoestrogens, and personal care products increased after digestion with both enzymes.

### Application to Breast Milk

The breast milk samples (n=15) from the short-term longitudinal breast milk study were investigated (Figure 3). For plasticizers, no clear trend was observed. For most of the phytoestrogens, BGS recombinant appeared to digest the conjugates at a higher level than *H. pomatia*. Only matairesinol and 8-prenylnaringenin resulted in higher concentration levels when treated with *H. pomatia* in comparison to BGS recombinant. The undigested and enzyme-treated samples showed similar results for the personal care products. In total, only five analytes were more efficiently digested with *H. pomatia*; whilst 14 analytes were cleaved at a higher rate with BGS recombinant. Here, only 35 xenobiotics out of the 85 measured xenobiotics were detected in the breast milk samples. This indicates that a low number of xenobiotics are actually transferred to the infant, in comparison to the 59 xenobiotics found in urine (Table S12).

In the long-term longitudinal breast milk study, where one mother collected her breast milk across a total of 87 days, 30 of the most contaminated samples according to Jamnik *et al*. were selected to compare the hydrolysis efficiencies of the two tested enzyme mixtures *H. pomatia* and BGS recombinant with an undigested control [24]. The majority of the signals were observed in three classes of xenobiotics, namely plasticizers, phytoestrogens, and personal care products.

For plasticizers, the concentration levels of only three analytes increased after hydrolysis, with *H. pomatia* displaying lower concentrations in two out of these three analytes most likely due to background contamination (Figure S1). BGS recombinant was able to produce on average higher concentration levels in mono-n-butyl-phthalate, and a slight increase in n-butylbenzenesulfonamide. For phytoestrogens, the samples treated with *H. pomatia* displayed higher concentration levels in seven analytes, in comparison to those treated with BGS recombinant, only displaying higher concentrations in three phytoestrogens. However, formononetin and resveratrol was only detected after hydrolysis with BGS recombinant. Finally, for the personal care products, four analytes were present at higher levels in the untreated samples. This can be traced back to the enzyme contamination in both enzymes. Overall, nine analytes were more efficiently cleaved with *H. pomatia*; whilst only five were more efficiently deconjugated when treated with BGS recombinant. From the 85 xenobiotics measured with our LC-MS/MS method, 34 analytes were detected in the breast milk samples from the mother, 18 of which only present after enzymatic treatment, thus indicating the presence of phase II metabolites in the breast milk samples (Table S11)

### Application to Plasma

To also include a human plasma sample, a commercial lithium-heparin pooled plasma sample was analyzed untreated and treated with *H. pomatia* and BGS recombinant enzymes in triplicate (9 samples). Both *H. pomatia* and BGS recombinant could hydrolyze a total of 24 xenobiotics (Table 1) Individual scrutiny of the xenobiotic classes, however, revealed that BGS recombinant was more efficient in digesting plasticizers. With the exception of bisphenol F (only detected in one of the samples digested with *H. pomatia* at a very low level of 0.46 ng/mL), BGS recombinant resulted in higher levels of the free contaminants compared to *H. pomatia*. In contrast, phytoestrogens and personal care products were generally more efficiently hydrolyzed by *H. pomatia* to consistently yield higher concentrations in 15 out of the total 21 detected phytoestrogens and personal care products. Daidzein, resveratrol, and xanthohumol, however, were only detected in the samples digested with BGS recombinant. Regarding digestion efficiencies, *H. pomatia* generally deconjugated more phase II biotransformation products, indicating a high efficiency in plasma samples. A total of 37 xenobiotics were identified in the plasma samples, 15 of which were only detected after enzymatic treatment. These data indicate that plasma is also an important matrix to monitor environmental exposure via phase II metabolites which are frequently not properly considered in this matrix.

**Table 1.** Results of the lithium-heparin plasma before and after enzymatic hydrolysis with the two selected enzymes from *H. pomatia* and BGS recombinant. Samples below the limit of quantitation are shown as <LOQ; whilst samples that were below the limit of detection are displayed as n.d. Results are given in ng/mL.

| Analytes in Plasma | Concentration [ng/mL] |  |  |  |  |  |
| --- | --- | --- | --- | --- | --- | --- |
|  | Untreated |  | Treated with *H. pomatia* |  | Treated with BGS |  |
|  | Mean | RSD [%] | Mean | RSD [%] | Mean | RSD [%] |
| **Plasticizer/Plastic Components** |  |  |  |  |  |  |
| Bisphenol A | 3.4 | 11 | 4.5 | 5.8 | 5.4 | 3.8 |
| Bisphenol F | n.d. | n.d. | 0.46 | 0.00 | n.d. | n.d. |
| Mono-n-butyl phthalate | 5.2 | 1.1 | 5.6 | 54 | 6.6 | 21 |
| N-butylbenzenesulfonamide | 4.0 | 2.7 | n.d. | n.d. | 2.5 | 4.6 |
| **Perfluorinated Alkylated Substances** |  |  |  |  |  |  |
| Perfluorooctanoic acid | 0.24 | 5.4 | 0.29 | 17 | 0.32 | 0.00 |
| **Industrial Side Products and Pesticides** |  |  |  |  |  |  |
| 2-naphthol | 4.0 | 4.7 | 0.24 | 9.8 | 2.1 | 20 |
| 4-octylphenol | n.d. | n.d. | 3.2 | 9.2 | 3.5 | 7.7 |
| 4-tert-octylphenol | 5.1 | 11 | 12 | 34 | 4.8 | 4.5 |
| Nonylphenol | 12 | 9.3 | 22 | 6.8 | 11 | 3.7 |
| **Endogenous Estrogens** |  |  |  |  |  |  |
| Estrone | 0.02 | 24 | 0.25 | 18 | 0.15 | 6.2 |
| **Phytoestrogens and Metabolites** |  |  |  |  |  |  |
| Coumestrol | n.d. | n.d. | 0.17 | 30 | n.d. | n.d. |
| Daidzein | 0.03 | 8.4 | n.d. | n.d. | 0.32 | 40 |
| Enterodiol | <LOQ | n.d. | 0.62 | 3.8 | 0.40 | 5.9 |
| Enterolactone | n.d. | n.d. | 9.7 | 7.8 | 5.3 | 7.0 |
| Equol | n.d. | n.d. | 0.49 | 8.3 | 0.36 | 2.8 |
| Formononetin | 0.002 | 13 | n.d. | n.d. | n.d. | n.d. |
| Genistein | 0.02 | 8.9 | 2.4 | 45 | 2.2 | 42. |
| Isoxanthohumol | n.d. | n.d. | n.d. | n.d. | <LOQ | n.d. |
| Matairesinol | n.d. | n.d. | 0.56 | 0.0 | 0.14 | 3.6 |
| Resveratrol | n.d. | n.d. | <LOQ | n.d. | 1.1 | 1.2 |
| **Mycoestrogens and Metabolites** |  |  |  |  |  |  |
| Alternariol monomethyl ether | <LOQ | n.d. | n.d. | n.d. | n.d. | n.d. |
| Zearalenone | n.d. | n.d. | 0.06 | 27 | n.d. | n.d. |
| **Personal Care Product Ingredients, Pharmaceuticals and Metabolites** |  |  |  |  |  |  |
| Benzophenone 1 | n.d. | n.d. | 0.04 | 11 | 0.02 | 11 |
| Benzophenone 2 | n.d. | n.d. | 0.04 | 4.7 | n.d. | n.d. |
| Benzylparaben | <LOQ | n.d. | 0.004 | 12 | 0.002 | 17 |
| Butylparaben | 0.01 | 0.0 | <LOQ | n.d. | n.d. | n.d. |
| Ethylparaben | 0.13 | 6.5 | 1.8 | 9.3 | 0.50 | 3.6 |
| Isobutylparaben | 0.01 | 1.4 | 0.02 | 14 | 0.02 | 0.0 |
| Methylparaben | 1.2 | 2.4 | 9.8 | 3.4 | 8.3 | 1.6 |
| Propylparaben | 0.12 | 6.1 | 1.6 | 9.8 | 0.73 | 3.6 |
| Triclosan | <LOQ | n.d. | 0.06 | 0.0 | 0.07 | 1.2 |
| **Air Pollutants** |  |  |  |  |  |  |
| Cotinine | 37.9 | 0.66 | 9.8 | 3.7 | 21 | 7.0 |
| trans-3-OH-cotinine | 23.4 | 0.57 | 9.7 | 2.2 | 23 | 6.1 |
| 1-OH-pyrene | n.d. | n.d. | 0.65 | 38 | 0.28 | 16.4 |
| 3-OH-phenanthrene | n.d. | n.d. | n.d. | n.d. | 0.08 | n.d. |

## Discussion

The most commonly used approach to include phase II metabolites of xenobiotics in the investigation of the total exposure is an enzymatic hydrolysis prior to analysis. This is necessary because conjugated standards are not easily accessible and are typically very expensive. One of the major enzymes utilized for hydrolysis is the β-glucuronidase and sulfatase from *H. pomatia*, due to its ability to deconjugate glucuronides and sulfates simultaneously [12, 14, 15]. Alternatively, commercially available recombinant enzyme preparations such as BGS (a mixture of β-glucuronidase and sulfatase) and ASPC arylsulfatase combined with *E. coli* glucuronidase, are available. Data on their ability to hydrolyse is limited to a small range of compounds.

A preliminary experiment to investigate the deconjugation ability of the selected enzymes was conducted (Figure 1). In plasma and urine, *H. pomatia* was the most efficient enzyme mixture for the majority of analytes. In breast milk, however, the enzyme mixtures, BGS recombinant and ASPC with *E. coli* typically cleaved the phase II analytes at a higher rate. Nevertheless, all three enzymes perfomed well for most analytes and the highest deconjugation efficiency depends on the analyte of interest despite the different enzyme activities used for deconjugation. A possible reason for these descripancies is the different isoforms of β-glucuronidases and arylsulfatases present in the crude *H. pomatia* enzyme mixture. Since recombinant enzymes might consist of a single or at least more limited number of enzyme isoforms than naturally present in, e.g. a gastrointestinal tract, higher nominal activities may even deconjugate less analytes than lower activities of the crude extract of gastric juices from *H. pomatia* [31]. This correlates to previous research conducted by Jain *et al*. (2021), where similar efficiencies for *H. pomatia* and ASPC arylsulfatase was observed [23]. Wakabayashi and Fishman (1960) compared multiple β-glucuronidases, including *H. pomatia* and *E. coli* on their activities and found that all enzymes performed similarly [32]. *E. coli* with ASPC mixture requires an additional sample preparation step and two enzymes, making it costly for no significant increase in efficiency compared to the other two tested enzyme mixtures (Table S8). For this reason, only *H. pomatia* and BGS recombinant were selected for further experiments.

However, *H. pomatia* has also been widely criticized because of contamination by various analytes due to the biological nature of *H. pomatia*. This could lead to potential false negative results after subtraction of the background signal, especially if the concentration in the sample is substantially lower than the level in the snail extract [12, 14, 33]. Hence, the enzyme is often purified if the appropriate equipment is available. Purification, however, requires suitable equipment and is a complex procedure [34–37]. During application to human biofluid samples, it was observed that the enzyme mixtures *H. pomatia* as well as BGS recombinant are contaminated with target analytes. *H. pomatia* has a high level of inherent background contamination for many plasticizers, phytoestrogens, and personal care products (Figure 2). BGS recombinant also generated signals for most of these analytes; albeit at lower levels compared to *H. pomatia*. This contamination may stem from plastic components or solutions used in the production of recombinant enzymes in cell culture and the extraction of the snail extract. In particular, phytoestrogens might originate from plant-based feed for *H. pomatia*. Contamination with similar compound classes has previously been described; however, the exact sources remain unknown [38, 39]. BGS recombinant has the advantage of reduced contamination levels compared to the natural *H. pomatia extract* therefore the detection of trace levels in human biomonitoring studies may be easier. In addition, the excellent deconjugation abilities of BGS recombinant for the majority of xenobiotics, further supports the application of this enzyme instead of the crude *H. pomatia* extract for biospecimen when certain toxicant classes are investigated.

In human biomonitoring, breast milk is an important biological matrix for investigating biomarkers of exposure as molecules in this biofluid can be potentially transferred to the infant. Toddlers are particularly susceptible to xenobiotic exposure as they are not yet fully developed [27, 40, 41]. Very few studies have investigated the presence of the detoxification products in breast milk. Phase II metabolites are generally considered to be less potent than the native compound, however, activation might also occur [8, 42]. In the long-term longitudinal breast milk study, plasticizers such as mono-n-butylphthalate and n-butylbenzenesulfonamide, and phytoestrogens such as formononetin and enterodiol were detected at higher concentration levels in the samples treated with BGS recombinant than those treated with *H. pomatia*. The highest concentration increase in certain breast milk samples observed after enzymatic treatment were from 4-tert-octylphenol (Table S11). When this environmental estrogen is transferred to the suckling child, it is of concern as it has been described to cause reproductive disorders in males such as low sperm counts, hypospadias, and cryptorchidism [43, 44]. Furthermore, formononetin levels also increased drastically after enzymatic treatment. To date, no traces of formononetin have been identified in breast milk as formononetin was primarily present in the conjugated form, deconjugation with *H. pomatia* was less efficient than the recombinant enzyme mixture and the heavy contamination of the crude *H. pomatia* extract might mask cleaved compounds. Thus, formononetin has not been described in previous studies using *H. pomatia* [45–47]. This phytoestrogen has a similar structure to 17-β-estradiol. Therefore, the substance can activate estrogen receptors by binding to the active site, interfering subsequently with the endocrine system [48, 49]. Furthermore, formononetin has shown potential as a chemotherapeutic drug by inhibiting tumour growth *in vivo* [50]. However, disruption of the hormonal homeostasis can also have detrimental effects, such as the induction of breast cancer [51]. In the short-term longitudinal breast milk study, one mother had very high levels of the plasticizer mono-2-ethylhexylphthalate, especially after enzymatic treatment in breast milk (Table S12). This plasticizer is a major component in flooring, clothing, packaging, and toys and can impact the development and reproductive health of infants [52, 53]. Further metabolites did not significantly increase after enzymatic hydrolysis.

In the long-term breast milk study, *H. pomatia* was able to hydrolyze nine analytes more efficiently than BGS recombinant, which could cleave only four xenobiotics at a higher rate (Table S11). However, in the short-term study, the results correlated more with the preliminary experiment including a different set of analytes, as BGS recombinant was able to digest 14 analytes more efficiently, in comparison to *H. pomatia*, only cleaving five xenobiotics more efficiently. Overall, when looking at both breast milk studies, compared to *H. pomatia*, BGS recombinant resulted in higher concentration levels and increased the number of detected analytes. Breast milk is a complex biofluid that is high in fat. Therefore, matrix effects might hamper the detection of low-concentrated analytes and the extraction efficiency is lower compared to more polar biofluids. Therefore, the lower contaminated BGS recombinant is potentially a better alternative to investigate phase II metabolites in breast milk compared to *H. pomatia* with high intrinsic concentrations of specific analytes.

In both breast milk studies phase II metabolites from plasticizers, pesticides, and phytoestrogens were present following enzymatic treatment of the samples. In general, very low concentration levels of free xenobiotics were found in the breast milk samples before hydrolysis, indicating a negligible amount of exposure to the infant. Although breast milk is very important for nourishing an infant, it may also be an excretion path of xenobiotics. Typically, breast milk substitutes and infant formulae contain higher contaminant levels and breast feeding should never be avoided or shortened because of (low-level) xenobiotic contamination [54]. Depending on the study, 34 or 35 xenobiotics were detected in breast milk, where 18 analytes were only detected after the enzymatic treatment, indicating that more than half of the analytes were in its conjugated form. The mother detoxified most of the xenobiotics before transporting them to her child, limiting exposure to the bioactive forms. Nevertheless, deconjugated analytes might be reactivated in the infant’s body. Despite its high lipid content, previous studies have shown that also polar phase II metabolites can be found in breast milk [16, 17].

The majority of metabolized xenobiotics are excreted through the urine. Therefore, the analysis of phase II conjugates is often conducted in this matrix [1, 8]. However, other matrices, such as breast milk also contain biotransformation products. In the short-term longitudinal breast milk study, not only breast milk, but also urine samples were collected by the same mothers. This enabled the comparison of the conjugation extent in both matrices. 59 xenobiotics were detected in urine samples compared to 35 in breast milk, thus most xenobiotics in the body were rather excreted through urine and only slightly more than half of them were transferred to the breast milk (Table S13). However, this comparison is not straight forward and needs to be interpreted in light of the specific physico-chemical and toxicokintic properties of a certain xenobiotic. *H. pomatia* could cleave most glucuronides and sulfates more efficiently in urine. However, both enzymes were able to digest the metabolites similarly, indicating that the enzymatic hydrolysis was successful with both enzymes (Figure 3). In this human biofluid, the inherent background contamination of *H. pomatia* did not lead to any large discrepancies in the results because overall the xenobiotics were present at higher levels. Hence, an additional purification step for this matrix is not really warranted, as it is time consuming, costly, and may introduce further contaminations due to additional preparation steps. High levels of 4-hydroxyestrone and enterolactone were observed in all three mothers after enzymatic treatment (Table S13). Although 4-hydroxyestrone has a protective effect against oxidative damage, the metabolite can also induce cancer proliferation [55, 56]. Nevertheless, enterolactone has been shown to inhibit ovarian cancer cell proliferation and the formation of metastases [57, 58].

Lithium-heparin pooled plasma samples were also investigated with *H. pomatia* and BGS recombinant for phase II metabolites. Several publications have indicated that phase II metabolites can be detected in plasma [12–15]. In this matrix, *H. pomatia* clearly hydrolyzed phase II conjugates more efficiently (Table 1). Similar to urine, the background contamination of *H. pomatia* had a lower impact on the detection of metabolites. Certain metabolite classes, such as plasticizers, however, were more efficiently digested with BGS recombinant. Both enzymes therefore confirmed the presence of phase II metabolites in plasma by verifying the metabolized state of xenobiotics in this commonly assessed biofluid of exposure.

Overall, in urine samples, most xenobiotics were detected (59 out of the 85 measured analytes) in comparison to breast milk where 34 and 35 analytes were measured, and plasma with 37 xenobiotics. Urine has a relatively high salt content and contains mostly polar compounds, as it consists mainly of urea, creatinine, proteins and inorganic salts. Breast milk, however, is a very complex matrix, containing high levels of lipids, proteins, and sugars; similarly to plasma, which additionally also contains inorganic salts and urea. These substances present in breast milk and plasma may interfere with measurements, especially the high sugar and fat contents might be troublesome. Signal suppression in the electrospray ionization by e.g. lactose source might be one reason for lower signal intensity in human breast milk [12, 59].

While very comprehensive in scope this study also has several limitations. The design was intended to cover different enzymes and three complex biological matrices that are by nature highly variable. Therefore, the results presented rather give a first global picture of de-conjugation capacities covering extremely diverse chemical classes than a final evaluation on which enzyme is the best solution for each matrix and application. Moreover, specific enzyme activities can be impacted by buffer capacities and the pH. The enzymes tested here were dissolved in different buffers (partially according to the suppliers information with proprietary information). When hydrolysing conjugates using H. pomatia, for a high number of analytes an interfering contamination was introduced to the sample that needed to be subtracted during data evaluation. This likely increased the measurement uncertainty. In addition, frequently the background signal was enhanced. It was not possible to fully align the enzyme activities since the ratio between β-glucuronidase and arylsulfatase activity in *H. pomatia* and BGS were fixed and the difference in enzyme activities were too high to allow for sufficient arylsulfatase activity when normalising to β-glucuronidase activity. Finally, it should be taken into account that the timing of breast milk and urine sampling impacts the exposure levels in the respective matrices and is not straight forward. It clearly needs to be considered in light of a molecules specific toxicokinetic properties.

## Conclusion and outlook

In conclusion, the most commonly utilized enzymes in the investigation of phase II conjugates, *H. pomatia*, remains in general the most efficient enzyme in urine and plasma. In breast milk, however, more analytes were detected with BGS recombinant; thus, providing promising results on the presence of phase II metabolites in this matrix. The recombinant enzyme treatment benefited from the lower background contamination. Nevertheless, BGS is the most cost intensive alternative because more enzyme for the deconjugation reactions is required to reach the same activity. The deconjugation depended highly on the matrix and compound class, therefore the optimal enzyme might vary between application. This study indicates *H. pomatia* as the preferable enzyme for urine and plasma samples, and BGS recombinant for breast milk samples. The hydrolysis step was kept simple, rapid, while providing high efficiency to be suitable in multi-class exposome-type biomonitoring and ExWAS studies. The enzymatic treatment increased the concentration levels of most deconjugated analytes with various structures revealing significant levels of phase II metabolites in urine, plasma and breast milk highlighting the transfer of masked xenobiotics to the infant. The investigation of multiple matrices concerning biotransformation products will be an essential step for better assessing the chemical exposome. While the integration of standards for conjugated xenobiotics might be desirable, their availability and complexity of biotransformation warrant the establishment of global deconjugation protocols. A possible solution for overcoming the issue of background contamination in *H. pomatia* is the purification of the raw enzyme extract that is commercially available by affinity chromatography as described by Ballet et al. [34]. However, this contradicts the high-throughput ambitions of current exposome research by adding an additional working step.

## Supporting information

SI_Part_A

SI_Part_B

## Acknowledgements

We would like to acknowledge all mothers who provided samples. Gratitude is also expressed to Ian Oesterle for his technical assistance and scientific input. We are also thankful to the Mass Spectrometry Centre (MSC) of the Faculty of Chemistry at the University of Vienna for their technical support. Camila Berner and Janet Jones (Kura Biotech) are acknowledged for providing enzymes and support in optimizing enzyme efficiencies. Finally, we highly appreciate Mario S.P. Correia from Uppsala University for his critical scientific input. We would also like to thank all group members of the Global Exposomics and Biomonitoring Laboratory at the University of Vienna for their support and assistance.

## Conflicts of interest

The authors have no relevant financial or non-financial interests to disclose.

## Notes

### Competing Interest Statement

The authors have declared no competing interest.

