## Supplementary material for "A broad, exposome-type evaluation of xenobiotic phase II biotransformation in human biofluids by LC-MS/MS": SI_Part_A

### Chemicals and Reagents

The following xenobiotic reference standards were purchased: 2-tert-butylphenol (2-tert-BP), 2-methoxyestradiol (2MeOE2), 2-methoxyestrone (2MeOE1), 2-naphthol, 3-benzylidencamphor (3-BC), 4-methylbenzyliden camphor (4-MBC), 4-octylphenol (4-OP), 4-tert-octylphenol (4-tert-OP), 5-hydroxymethylfurfural (HMF), 8-prenylnaringenin (8-Pn), α-zearalanol (α-ZAL), alternariol, anisodamine, benzophenone 1, benzophenone 2, benzyl butyl phthalate, benzylparaben, β-zearalanol (β-ZAL), bisphenol A (BPA), bisphenol AF (BPAF), bisphenol B (BPB), bisphenol C (BPC), bisphenol F (BPF), bisphenol S (BPS), butylparaben, chloroacetic acid, cotinine, daidzein, dibromoacetic acid, dibutyl phthalate, dichloroacetic acid, enterodiol, enterolactone, estradiol (E2), estriol (E3), estrone (E1), ethylparaben, fenarimol, formononetin, isobutylparaben, matairesinol, methiocarb, methylparaben, mono-2-ethylhexyl phthalate (MEHP), mono-n-butyl phthalate (MBP), N-butylbenzenesulfonamide, N-nitrosodimethylamine (NDMA), nonylphenol, octyl methoxycinnamate (OMC), perfluorooctanoic acid (PFOA), perfluorooctanesulfonic acid (PFOS), p-hydroxybenzoic acid (pOHBA), prochloraz, propylparaben, resveratrol, tetrabromobisphenol A (TBPA), triclosan, and zearalanone (ZAN) were obtained from Sigma-Aldrich (Vienna, Austria). $^{13}C_{18}$-zearalenone (ZEN), β-zearalenol (β-ZEL), and zearalenone (ZEN) were purchased from RomerLabs (Tulln, Austria). 16-epiestriol (16EpiE3), 16-α-hydroxyestrone (16OHE1), 17-epiestriol (17EpiE3), 1-hydroxypyrene, 2-hydroxyestradiol (2OHE2), 3-hydroxybenzo[a]pyrene (3-OH-BaP), 3-hydroxyphenanthrene, 4-hydroxyestrone (4OHE1), 4-methoxyestradiol (4MeOE2), 4-methoxyestrone (MeOE1), 5-hydroxymethyl-2-furanoic acid (HMFA), alternariol monomethyl ether (AME), acrylamide, aristolactam I, aristolochic acid I, bromoacetic acid, ethinylestradiol, glycidamide, glycitein, N-nitrosodiethylamine (NDEA), PhIP, scopolamine, and trans-3-hydroxy-cotinine were purchased from Toronto Research Chemicals (Toronto, Canada). Coumestrol was obtained from Enzo Life Sciences (New York, USA), whilst d4-genistein was purchased from CDN isotopes (Pointe-Claire, Canada), equol from Cayman Chemical Company (Michigan, USA), jacobine, jacobine-N-oxide, riddelliin, riddelliin-N-oxide from PhytoPlan (Heidelberg, Germany), and xanthohumol from Extrasynthese (Genay, France). Zearalenone-14-glucuronide (ZEN-14-GlcA) was synthesised at the Technical University of Vienna and generously provided by Prof. Mikula, Zearalenone-14-sulfate (ZEN-14-sulfate) was kindly provided by Prof. Berthiller from the University of Natural Resources and Life Sciences, Vienna (BOKU).

**Table S 1.** LC-MS/MS transitions and parameters of phytoestrogens and synthetic chemicals added to the method from Jamnik et al. (2022) Preliminary experiment testing the efficiencies of the three enzyme mixtures H. pomatia, BGS recombinant, and E. coli with ASPC recombinant [1].

| Compound | MS/MS Transitions | | MS/MS parameters | | | | | |
|---|---|---|---|---|---|---|---|---|
| | Q1 Mass [Da] | Q3 Mass [Da] | Polarity | Retention Time [min] | DP [volts] | EP [volts] | CE [volts] | CXP [volts] |
| **Phytoestrogens and Metabolites** | | | | | | | | |
| Genistein 7-sulfate | 348.798 | 268.900 | neg | 4.10 | -75.000 | -10.000 | -38.000 | -53.000 |
| | | 133.100 | | | | | -58.000 | -7.000 |
| Caffeic acid 3-β-D-glucuronide | 355.074 | 135.000 | neg | 0.65 | -90.000 | -10.000 | -36.000 | -21.000 |
| | | 179.200 | | | | | -30.000 | -5.000 |
| Daidzein 7-β-D-glucuronide | 429.141 | 174.700 | neg | 2.79 | -50.000 | -10.000 | -16.000 | -27.000 |
| | | 113.000 | | | | | -22.000 | -17.000 |
| Genistein 7-β-D-glucuronide | 444.932 | 113.100 | neg | 3.10 | -45.000 | -10.000 | -24.000 | -19.000 |
| | | 269.000 | | | | | -40.000 | -45.000 |
| Kaempferol 3-O-glucuronide | 460.963 | 284.900 | neg | 3.30 | -40.000 | -10.000 | -32.000 | -17.000 |
| | | 112.900 | | | | | -20.000 | -17.000 |
| Quercetin 7-O-β-D-glucuronide | 477.062 | 300.800 | neg | 3.00 | -5.000 | -10.000 | -28.000 | -25.000 |
| | | 150.900 | | | | | -48.000 | -11.000 |
| **Synthetic Chemicals** | | | | | | | | |
| 4-nitrocatechol-sulfate | 234.000 | 154.000 | neg | 3.39 | -60.000 | -10.000 | -20.000 | -11.000 |
| | | 124.000 | | | | | -38.000 | -11.000 |
| Phenolphthalein β-D-glucuronide | 493.000 | 93.000 | neg | 4.06 | -60.000 | -10.000 | -40.000 | -11.000 |
| | | 175.000 | | | | | -30.000 | -11.000 |
| Phenolphthalein | 317.100 | 273.100 | neg | 0 – end of retention time | -60.000 | -10.000 | -25.000 | -11.000 |
| | | 93.000 | | | | | -40.000 | -11.000 |
| Nitrocatechol | 154.000 | 124.000 | neg | 0 – end of retention time | -60.000 | -10.000 | -25.000 | -11.000 |
| | | 95.000 | | | | | -35.000 | -11.000 |
| p-cresol-sulfate | 107.100 | 105.900 | neg | 0 – end of retention time | -80.000 | -10.000 | -23.000 | -12.000 |
| | | 92.000 | | | | | -32.000 | -12.000 |

**Table S 2.** Calibration range, quantification limit (LOQ), detection limit (LOD), and extraction recovery ($R_E$) results of the method from Jamnik et al. (2022). The values are provided as concentration in ng/mL in urine (U), plasma (P), and breast milk (M). Parameters which were unable to be determined are given as n.d.

| Compound | Calibration range [ng/mL] | LOQ [ng/mL] | | | LOD [ng/mL] | | | Extraction recovery [$R_E$] | | |
|---|---|---|---|---|---|---|---|---|---|---|
| | U/P/M | U | P | M | U | P | M | U | P | M |
| **Plasticizer/Plastic Components** | | | | | | | | | | |
| Bisphenol A | 0.03-10/0.1-10/0.1-10 | 0.21 | 0.57 | 0.40 | 0.06 | 0.17 | 0.12 | 1.16 | 1.04 | 0.82 |
| Bisphenol AF | 0.015-5 | 0.04 | 0.08 | 0.04 | 0.01 | 0.02 | 0.01 | 0.97 | 0.96 | 0.47 |
| Bisphenol B | 0.03-1 | 0.02 | 0.04 | 0.03 | 0.01 | 0.01 | 0.01 | 0.91 | 0.76 | 0.53 |
| Bisphenol C | 0.06-20/0.2-20/0.2-20 | 0.22 | 0.85 | 0.24 | 0.07 | 0.26 | 0.07 | 0.95 | 0.82 | 0.5 |
| Bisphenol F | 0.05-5 | 0.06 | 0.07 | 0.03 | 0.02 | 0.02 | 0.01 | n.d. | 0.73 | 0.72 |
| Bisphenol S | 0.002-0.2 | 0.04 | 0.01 | 0.01 | 0.01 | 0.00 | 0.00 | 0.87 | 1.09 | 0.95 |
| Mono-n-butyl phthalate | 0.15-50 | 0.34 | 0.20 | 0.16 | 0.10 | 0.06 | 0.05 | 0.95 | 1.38 | 0.78 |
| Mono-2-ethylhexyl phthalate | 0.045-15/n.d./0.045-4.5 | 0.05 | n.d. | 3.80 | 0.01 | n.d. | 1.14 | n.d. | n.d. | 0.47 |
| N-butylbenzenesulfonamide | 0.3-100 | 1.20 | 1.00 | 1.50 | 0.36 | 0.30 | 0.45 | 0.96 | 0.87 | 0.71 |
| Benzylbutylphthalate | 0.075-25/0.075-25/n.d. | 0.18 | 0.66 | n.d. | 0.05 | 0.20 | n.d. | 0.8 | 0.78 | n.d. |
| Dibutylphthalate | 1.5-500/1.5-500/n.d. | 3.10 | 7.60 | n.d. | 0.93 | 2.28 | n.d. | 0.36 | 0.95 | n.d. |
| Tetrabrombisphenol A | 0.03-10/0.3-10/n.d. | 0.02 | 0.33 | n.d. | 0.01 | n.d. | n.d. | 0.91 | 0.56 | n.d. |
| **Perfluorinated Alkylated Substances** | | | | | | | | | | |
| Perfluoroctanoic acid | 0.015-1.5/0.0045-1.5/0.0045-0.45 | 0.22 | 0.18 | 0.09 | 0.07 | 0.05 | 0.03 | 1.19 | 0.84 | 0.75 |
| Perfluoroctanesulfonic acid | 0.045-1.5/0.015-1.5/0.015-1.5 | 0.05 | 0.14 | 0.02 | 0.01 | 0.04 | 0.00 | 1 | n.d. | 0.95 |
| **Industrial Side Products and Pesticides** | | | | | | | | | | |
| 2-naphthol | 0.009-3/0.01-3/0.009-3 | 0.02 | 0.03 | 0.02 | 0.01 | 0.01 | 0.01 | 0.95 | 0.96 | 0.76 |
| Methiocarb | 0.05-0.5/0.015-0.5/0.015-0.5 | 0.02 | 0.04 | 0.02 | 0.01 | 0.01 | 0.00 | n.d. | n.d. | 0.38 |
| Prochloraz | 0.0005-0.05/0.0015-0.05/0.015-0.05 | 0.00 | 0.00 | 0.05 | 0.00 | 0.00 | 0.02 | n.d. | n.d. | n.d. |
| 2-tert-Butylphenol | 15-5000 | 34.00 | 63.00 | 98.00 | 10.20 | 18.90 | 29.40 | n.d. | 0.05 | 0.2 |
| 4-octylphenol | 3-300/3-1000/30-1000 | 1.40 | 2.30 | 24.00 | 0.42 | 0.69 | 7.20 | 0.43 | 1.06 | n.d. |
| 4-tert-octylphenol | 0.45-150/0.45-150/15-45 | 0.14 | 3.30 | 40.00 | 0.04 | 0.99 | 12.00 | n.d. | 0.8 | n.d. |
| Fenarimol | 0.003-0.3/0.003-3/0.009-0.3 | 0.00 | 0.01 | 0.01 | 0.00 | 0.00 | 0.00 | 0.88 | 0.95 | n.d. |
| Nonylphenol | 0.75-250/7.5-250/n.d. | 3.20 | 1.60 | n.d. | 0.96 | 0.48 | n.d. | 0.37 | 1.21 | n.d. |
| **Endogenous Estrogens** | | | | | | | | | | |
| Estrone | 0.003-0.3/0.009-0.3/0.009-0.3 | 0.01 | 0.01 | 0.01 | 0.00 | 0.00 | 0.00 | 1.29 | 0.82 | n.d. |
| Estradiol | 0.09-3 | 0.05 | 0.08 | 0.09 | 0.02 | 0.02 | 0.03 | 1.07 | 0.76 | n.d. |
| Estradiol-17-glucuronide | n.d./1.5-5/0.15-5 | n.d. | 4.90 | 0.91 | n.d. | 1.47 | 0.27 | n.d. | n.d. | 0.77 |
| Estradiol-3-sulfate | 0.045-1.5/0.015-1.5/0.0045-1.5 | 0.20 | 0.01 | 0.01 | 0.06 | 0.00 | 0.00 | n.d. | 0.77 | 0.56 |
| Estriol | 0.09-3/0.03-3/0.09-3 | 0.12 | 0.06 | 0.11 | 0.04 | 0.02 | 0.03 | 0.95 | 1.04 | 0.6 |
| 16-epiestriol | 0.3-10 | 0.23 | 0.13 | 0.19 | 0.07 | 0.04 | 0.06 | 0.91 | 0.93 | 0.57 |
| 16-α-hydroxyestrone | 0.045-1.5 | 0.10 | 0.03 | 0.04 | 0.03 | 0.01 | 0.01 | 0.67 | 0.82 | 0.74 |
| 17-epiestriol | 0.03-10/0.1-10/0.1-10 | 0.12 | 0.09 | 0.11 | 0.04 | 0.03 | 0.03 | 0.94 | 0.86 | 0.57 |

| Compound | Calibration range [ng/mL] | LOQ [ng/mL] | | | LOD [ng/mL] | | | Extraction recovery [$R_E$] | | |
|---|---|---|---|---|---|---|---|---|---|---|
| | U/P/M | U | P | M | U | P | M | U | P | M |
| 2-methoxy estrone | 0.025-2.5/0.075-2.5/0.075-2.5 | 0.05 | 0.05 | 0.05 | 0.01 | 0.02 | 0.01 | 0.97 | 0.97 | 0.43 |
| 2-methoxy estradiol | 0.006-2/0.02-2/0.06-2 | 0.02 | 0.03 | 0.03 | 0.01 | 0.01 | 0.01 | 0.99 | 0.9 | n.d. |
| 4-methoxy estrone | 0.005-0.5/0.015-0.5/0.015-0.5 | 0.02 | 0.01 | 0.01 | 0.00 | 0.00 | 0.00 | 0.96 | 0.83 | n.d. |
| 4-methoxy estradiol | 0.01-1/0.03-1/0.03-1 | 0.02 | 0.02 | 0.01 | 0.01 | 0.00 | 0.00 | 0.95 | 0.91 | 0.27 |
| 4-hydroxy estrone | 0.005-0.5/0.15-5/0.015-0.5 | 0.01 | n.d. | 0.01 | 0.00 | n.d. | 0.00 | 0.79 | n.d. | n.d. |
| **Phytoestrogens and Metabolites** | | | | | | | | | | |
| 8-prenylnaringenin | 0.009-3/0.009-3/0.03-3 | 0.01 | 0.02 | 0.02 | 0.00 | 0.01 | 0.00 | 0.88 | 0.87 | 0.39 |
| Coumestrol | 0.005-0.5/0.015-0.5/0.015-0.5 | 0.01 | 0.01 | 0.00 | 0.00 | 0.00 | 0.00 | 0.96 | 0.94 | 0.58 |
| Daidzein | 0.005-0.5/0.005-0.5/0.0015-0.5 | 0.03 | 0.02 | 0.01 | 0.01 | 0.00 | 0.00 | 0.98 | 0.84 | n.d. |
| Enterodiol | 0.015-0.5/0.005-0.5/0.0015-0.5 | 0.18 | 0.00 | 0.00 | 0.05 | 0.00 | 0.00 | 0.89 | 0.92 | 0.43 |
| Enterolactone | 0.06-20 | 1.10 | 0.17 | 0.17 | 0.33 | 0.05 | 0.05 | 1.01 | 0.9 | 0.77 |
| Equol | 0.006-2/0.02-2/0.02-2 | 0.01 | 0.03 | 0.01 | 0.00 | 0.01 | 0.00 | 0.97 | 0.97 | 0.6 |
| Formononetin | 0.00075-0.25 | 0.00 | 0.00 | 0.00 | 0.00 | 0.00 | 0.00 | n.d. | 0.94 | 0.58 |
| Genistein | 0.005-0.5/0.015-0.5/0.005-0.5 | 0.01 | 0.02 | 0.02 | 0.00 | 0.01 | 0.01 | 1.07 | 0.71 | 1.84 |
| Glycitein | 0.015-5/0.05-5/0.0015-0.5 | 0.13 | 0.06 | 0.00 | 0.04 | 0.02 | 0.00 | n.d. | 1.07 | 0.72 |
| Isoxanthohumol | 0.001-0.1/0.003-0.1/0.001-0.1 | 0.01 | 0.01 | 0.01 | 0.00 | 0.00 | 0.00 | 0.91 | 0.83 | n.d. |
| Matairesinol | 0.05-5/0.05-5/0.15-5 | 0.65 | 0.11 | 0.13 | 0.20 | 0.03 | 0.04 | 0.78 | 0.83 | 0.83 |
| Resveratrol | 1.5-150/0.45-45/0.45-150 | 2.20 | 0.92 | 0.30 | 0.66 | 0.28 | 0.09 | 0.93 | 0.72 | 0.1 |
| Xanthohumol | 0.03-10/0.1-10/0.1-3 | 0.05 | 0.14 | 0.22 | 0.02 | 0.04 | 0.07 | 0.91 | 0.83 | n.d. |
| **Mycoestrogens and Metabolites** | | | | | | | | | | |
| Alternariol | 0,1-10 | 0.16 | 0.14 | 0.08 | 0.05 | 0.04 | 0.02 | 0.94 | 0.91 | 0.61 |
| Alternariol monomethyl ether | 0.015-0.5 | 0.01 | 0.01 | 0.02 | 0.00 | 0.00 | 0.01 | 0.94 | 0.66 | n.d. |
| α-zearalanol | 0.05-5/0.015-5/0.05-5 | 0.10 | 0.07 | 0.09 | 0.03 | 0.02 | 0.03 | 0.97 | 0.96 | 0.44 |
| β-zearalanol | 0.015-5/0.05-5/0.05-5 | 0.13 | 0.06 | 0.05 | 0.04 | 0.02 | 0.02 | 1.02 | 0.91 | 0.54 |
| α-zearalenol | 0.02-0.2/0.02-0.2/0.06-0.2 | 0.01 | 0.01 | 0.06 | 0.00 | 0.00 | 0.02 | n.d. | n.d. | n.d. |
| β-zearalenol | 0.1-10 | 0.40 | 0.24 | 0.16 | 0.12 | 0.07 | 0.05 | 1 | 0.94 | 0.51 |
| α-zearalenol-14-glucuronide | n.d./0.45-1.5/0.045-1.5 | n.d. | 0.66 | 0.02 | n.d. | 0.20 | 0.01 | n.d. | n.d. | 0.61 |
| β-zearalenol-14-glucuronide | n.d./n.d./0.045-1.5 | n.d. | 0.42 | 0.04 | n.d. | 0.13 | 0.01 | n.d. | n.d. | 0.64 |
| Zearalanone | 0.09-3 | 0.08 | 0.20 | 0.12 | 0.02 | 0.06 | 0.04 | 0.98 | 0.88 | n.d. |
| Zearalenone | 0.009-3/0.03-3/0.09-3 | 0.03 | 0.03 | 0.09 | 0.01 | 0.01 | 0.03 | 0.98 | 0.86 | n.d. |
| Zearalenone-14-glucuronide | n.d./0.5-5/0.15-5 | n.d. | 1.20 | 0.09 | n.d. | 0.36 | 0.03 | n.d. | n.d. | 0.68 |
| Zearalenone-14-sulfate | 0.015-1.5/0.015-1.5/0.0045-1.5 | 0.15 | 0.02 | 0.02 | 0.05 | 0.01 | 0.00 | 0.99 | 0.91 | 0.46 |
| **Personal Care Product Ingredients, Pharmaceuticals and Metabolites** | | | | | | | | | | |
| Benzophenone 1 | 0.006-2 | 0.01 | 0.02 | 0.01 | 0.00 | 0.01 | 0.00 | 0.98 | 0.96 | 0.58 |
| Benzophenone 2 | 0.015-1.5/0.0045-1.5/0.0045-1.5 | 0.02 | 0.03 | 0.01 | 0.01 | 0.01 | 0.00 | 0.97 | 0.89 | 0.75 |
| Benzylparaben | 0.0045-0.15 | 0.00 | 0.00 | 0.00 | 0.00 | 0.00 | 0.00 | 0.97 | 0.92 | 0.51 |

| Compound | Calibration range [ng/mL] U/P/M | LOQ [ng/mL] U | P | M | LOD [ng/mL] U | P | M | Extraction recovery [$R_E$] U | P | M |
|---|---|---|---|---|---|---|---|---|---|---|
| Butylparaben | 0.003-1/0.01-1/0.003-1 | 0.01 | 0.01 | 0.01 | 0.00 | 0.00 | 0.00 | 0.87 | 0.85 | 0.47 |
| Ethylparaben | 0.003-1/0.003-1/0.01-1 | 0.00 | 0.02 | 0.04 | 0.00 | 0.01 | 0.01 | 0.93 | 0.81 | 1.03 |
| Isobutylparaben | 0.01-1/0.01-1/0.003-1 | 0.01 | 0.01 | 0.01 | 0.00 | 0.00 | 0.00 | 0.87 | 0.91 | 0.45 |
| Methylparaben | 0.0075-2.5 | 0.05 | 0.04 | 0.03 | 0.01 | 0.01 | 0.01 | 0.95 | 0.74 | 0.85 |
| Propylparaben | 0.006-2 | 0.01 | 0.01 | 0.01 | 0.00 | 0.00 | 0.00 | 0.93 | 0.8 | 0.78 |
| Ethinylestradiol | 0.1-10/0.1-10/0.3-10 | 0.04 | 0.12 | 0.21 | 0.01 | 0.04 | 0.06 | 0.97 | 0.93 | n.d. |
| 3-benzylidencamphor | 4.5-450/4.5-1500/4.5-450 | 0.30 | 2.00 | 6.10 | 0.09 | 0.60 | 1.83 | n.d. | 0.34 | n.d. |
| 4-methylbenzylidencamphor | 0.45-150/0.45-150/1.5-150 | 0.43 | 0.41 | 14.00 | 0.13 | 0.12 | 4.20 | n.d. | 0.64 | n.d. |
| Octyl methoxycinnamate | 200-600/60-200; 200-600; 600-2000/n.d. | 160.00 | 260.00 | 730.00 | 48.00 | 78.00 | 219.00 | n.d. | n.d. | n.d. |
| p-hydroxybenzoic acid | n.d./5-500/1.5-500 | n.d. | 9.80 | 7.60 | n.d. | 2.94 | 2.28 | n.d. | 0.76 | 0.74 |
| Triclosan | 0.03-10/0.03-10/0.3-10 | 0.04 | 0.05 | 1.00 | 0.01 | 0.01 | 0.30 | 0.72 | 1 | n.d. |
| **Phytotoxins** | | | | | | | | | | |
| Anisodamine | 0.05-0.5/0.005-0.5/0.0015-0.5 | 0.07 | 0.01 | 0.00 | 0.02 | 0.00 | 0.00 | n.d. | 1.06 | 0.43 |
| Aristolochic acid I | 0.3-10/0.3-10/0.1-10 | 0.33 | 0.24 | 0.44 | 0.10 | 0.07 | 0.13 | 0.78 | 0.73 | n.d. |
| Aristolactam I | 0.015-5/0.05-5/0.15-5 | 0.02 | 0.02 | 0.20 | 0.00 | 0.01 | 0.06 | 0.93 | 0.84 | n.d. |
| Jacobine | 0.075-2.5/0.075-2.5/0.025-2.5 | 0.40 | 0.04 | 0.04 | 0.12 | 0.01 | 0.01 | n.d. | 0.9 | 0.45 |
| Jacobine-N-oxide | 0.05-0.5/0.015-0.5/0.0015-0.5 | 0.06 | 0.01 | 0.00 | 0.02 | 0.00 | 0.00 | n.d. | 1.02 | 0.29 |
| Riddelliin | 0.3-3/0.09-3/0.03-3 | 1.20 | 0.18 | 0.12 | 0.36 | 0.05 | 0.04 | n.d. | 0.88 | 0.43 |
| Riddelliin-N-oxide | 0.6-2/0.06-2/0.006-2 | 1.00 | 0.04 | 0.02 | 0.30 | 0.01 | 0.01 | n.d. | 1.01 | 0.39 |
| Scopolamine | 0.00045-0.15 | 0.00 | 0.00 | 0.00 | 0.00 | 0.00 | 0.00 | 1.02 | 0.96 | 0.56 |
| **Disinfection By-Products** | | | | | | | | | | |
| Bromoacetic acid | n.d./2-200/n.d. | n.d. | 79.00 | n.d. | n.d. | 23.70 | n.d. | n.d. | n.d. | n.d. |
| Dibromoacetic acid | 4.5-150/0.45-150/0.45-150 | 32.00 | 1.80 | 2.70 | 9.60 | 0.54 | 0.81 | n.d. | 0.83 | 0.37 |
| Dichloroacetic acid | 22.5-750/2.25-750/7.5-750 | 60.00 | 10.00 | 6.60 | 18.00 | 3.00 | 1.98 | n.d. | 1.01 | 0.29 |
| **Food Processing By-Products** | | | | | | | | | | |
| Acrylamide | 30-1000/10-1000/10-1000 | 92.00 | 5.30 | 22.00 | 27.60 | 1.59 | 6.60 | n.d. | 0.64 | 0.28 |
| 5-hydroxymethylfurfural | 3.5-350/1.05-350/1.05-350 | 29.00 | 16.00 | 5.60 | 8.70 | 4.80 | 1.68 | n.d. | 0.75 | n.d. |
| 5-hydroxymethyl-2-furanoic acid | n.d./60-2000/6-2000 | n.d. | 150.00 | 22.00 | n.d. | 45.00 | 6.60 | n.d. | 0.86 | 0.5 |
| N-nitrosodimethylamine | 30-3000/90-3000/30-3000 | 240.00 | 230.00 | 170.00 | 72.00 | 69.00 | 51.00 | n.d. | n.d. | n.d. |
| PhIP | 0.003-1/0.01-1/0.003-1 | 0.01 | 0.01 | 0.00 | 0.00 | 0.00 | 0.00 | 0.96 | 0.93 | 0.55 |
| **Air Pollutants** | | | | | | | | | | |
| Cotinine | 0.045-15/0.45-15/0.045-15 | 0.89 | 0.05 | 0.11 | 0.27 | 0.02 | 0.03 | 0.86 | n.d. | n.d. |
| Trans-3-hydroxy-cotinine | 0.15-5/0.05-5/0.015-5 | 2.30 | 0.07 | 0.03 | 0.69 | 0.02 | 0.01 | n.d. | n.d. | n.d. |
| 1-hydroxy-pyrene | 0.06-20/0.2-20/0.2-6 | 0.19 | 0.22 | 1.30 | 0.06 | 0.07 | 0.39 | 0.91 | 0.62 | 1.04 |
| 3-hydroxy-phenanthrene | 0.015-1.5/0.045-1.5/0.045-1.5 | 0.02 | 0.03 | 0.05 | 0.01 | 0.01 | 0.02 | 0.92 | 0.87 | n.d. |

**Table S 3.** Concentration of reference standards in the sample and multi-analyte mix in ng/mL for the reference glucuronide (GlcA) and sulfate (Sulf) conjugates.

| Analytes | Conc. in sample [ng/mL] | Conc. of mix [ng/mL] |
|---|---|---|
| Kaempferol 3-O-GlcA | 5 | 25 |
| Quercetin 7-O-B-D-GlcA | 12 | 60 |
| Genistein 7-B-D-GlcA | 8 | 40 |
| Caffeic Acid 3-B-D-GlcA | 10 | 50 |
| Daidzein 7-B-D-GlcA | 3 | 15 |
| Genistein 7-Sulf | 1.5 | 7.5 |
| Estradiol-17-GlcA | 0.5 | 2.5 |
| Estradiol-3-Sulf | 1 | 5 |
| p-Cresol-Sulf | 4 | 20 |
| Phenolpht-GlcA | 0.15 | 0.75 |
| Nitrocatechol-Sulf | 0.03 | 0.15 |

**Table S 4.** Activity of the three tested enzyme mixtures H. pomatia, BGS, and ASPC with E. coli based on the activities obtained from the enzyme assays as in the pure enzyme, in enzyme and buffer solution and in the sample.

| Type of solution | Activity [U/mL] | *H. pomatia* | BGS | ASPC / *E. coli* |
|---|---|---|---|---|
| **Pure enzyme** | **β-glucuronidase** | 27082 | 638109 | 472656 |
| | **arylsulfatase** | 141.49 | 18.08 | 23.37 |
| **In enzyme solution** | **β-glucuronidase** | 4000 | 638109 | 472656 |
| | **arylsulfatase** | 49 | 18.08 | 23.37 |
| **In sample** | **β-glucuronidase** | 2000 | 212703 | 2000 |
| | **arylsulfatase** | 24.5 | 7.8 | 6.03 |

### Optimisation of pH

To determine the amount of buffer required to provide optimal conditions for enzyme activity, the pH values of 100 µL of pooled breast milk, urine, or citrate plasma was tested with a sensitive pH paper (Supelco, Merck 1.09542.0001) with a range of pH 4.0–7.0 ± 0.2–0.5 before and after the addition of buffer. Different volumes (50 µL, 100 µL, and 120 µL) of the buffers were tested. The optimal pH-value for BGS recombinant, *E. coli* and ASPC recombinant was neutral and PBS, instant buffer II, and ammonium carbonate buffers were tested (refer to Table S3). For *H. pomatia* the optimal pH range of 4.5 – 5.0 was tested with ammonium acetate buffer. The volume with a pH closest to the optimal hydrolysis conditions for the various enzymes was used in the further sample preparation.

To optimize conditions and create a favourable environment to achieve the highest enzyme efficiency, the pH of the three matrices, urine, plasma, and breast milk were tested before and after the addition of various buffers. As shown in supplementary Table S4, it was found that an equivalent volume of buffer to volume of matrix was required to provide a pH range closest to the optimal pH value (*i.e.,* 100 µL buffer with 100 µL matrix or 50 µL buffer with 50 µL matrix) for all enzyme buffer combinations. As phosphates inhibit arylsulfatase activities (present in PBS buffer) and ammonium carbonate buffer is only stable for a short period of time, the instant buffer II was used in further experiments for enzymes requiring neutral environments [2].

**Table S 5.** Average pH of the tested matrices, where the average of the samples used for matrix matched standards were tested in comparison with the optimal pH values of the tested enzyme mixtures.

| **Matrix** | **Average matrix pH** | **Optimal pH for enzyme** | | |
|---|---|---|---|---|
| | | ***H. pomatia*** | **BGS** | ***E. coli* / ASPC** |
| **Urine** | 6.4 | | | |
| **Plasma** | 8 | 4.5 - 5.0 | 6.8 -7.5 | 6.8 - 7.5 |
| **Breast Milk** | 6.7 | | | |

**Table S 6.** Determination of the amount of buffer needed to achieve the optimal pH values of the tested enzyme mixtures. The pH values were measures after the addition of (i) 50 µL, (ii) 100 µL, and (iii) 120 µL of the three buffers to 100 µL of matrix.

| Matrix | Buffer | Amount of buffer added to 100 µL matrix | | |
|---|---|---|---|---|
| | | 50 µL | 100 µL | 120 µL |
| Urine | **2.5 M ammonium acetate** | 5.3 | 5.1 | 5.1 |
| | **160 mM ammonium carbonate** | 6.7 | 6.6 | 6.5 |
| | **PBS** | 6.4 - 6.5 | 6.7 | 6.7 |
| Plasma | **2.5 M ammonium acetate** | 5.8 | 5.5 | 5.3 |
| | **160 mM ammonium carbonate** | 7 | 6.7 - 6.8 | 6.6 - 6.7 |
| | **PBS** | 8 | 7.4 - 7.3 | 7 |
| Breast Milk | **2.5 M ammonium acetate** | 5.8 | 5.5 | 5.3 |
| | **160 mM ammonium carbonate** | 6.7 | 6.8 - 7.0 | 7 |
| | **PBS** | 6.4 | 6.5 - 6.6 | 6.7 |

**Table S 7** Activities of the enzyme mixtures compared within this work. β-glucuronidase activity is based on the ability to deconjugate phenolphthalein β-glucuronidase and the sulfatase activity is provided according to the hydrolysis efficiency of p-nitrocatechol-sulfate and is provided in units per millilitres.

| Enzyme | Activity in U/mL | | |
|---|---|---|---|
| | BGS | *E. coli* / ASPC | *H. pomatia* |
| β-glucuronidase | 644118 | 476777 | 27363 |
| Arylsulfatase (initial) | 0.75 | 0.88 | 656 |
| Arylsulfatase (repeat) | 18.1 | 23.4 | - |

**Table S 8.** Student t-test comparison between the hydrolysis efficiencies of the three tested enzyme mixtures. The student t-test compared the data of H. pomatia against BGS recombinant, H. pomatia against E. coli with ASPC recombinant, and BGS recombinant against E. coli with ASPC recombinant. p values below 0.05 were considered significant (n.s. $p > 0.05$, * $p < 0.05$, ** $p < 0.01$, *** $p < 0.001$).

| Analytes | *H. pomatia* - BGS | *H. pomatia* - *E. coli* / ASPC | BGS - *E. coli* / ASPC |
|---|---|---|---|
| | Urine | | |
| Kaempferol | ** | ** | n.s. |
| Quercetin | n.s. | n.s. | n.s. |
| Genistein | ** | ** | n.s. |
| Caffeic Acid | * | * | n.s. |
| Daidzein | ** | * | *** |
| Estradiol | n.s. | n.s. | * |
| Cresol | *** | n.s. | *** |
| Nitrocatechol | *** | *** | ** |
| Phenolphthalein | n.s. | ** | ** |
| **Analytes** | **Plasma** | | |
| Kaempferol | *** | *** | n.s. |
| Quercetin | n.s. | * | n.s. |
| Genistein | * | * | n.s. |
| Caffeic Acid | *** | *** | ** |
| Daidzein | *** | *** | *** |
| Estradiol | *** | ** | * |
| Cresol | *** | *** | *** |
| Nitrocatechol | *** | *** | *** |
| Phenolphthalein | * | * | n.s. |
| **Analytes** | **Breast Milk** | | |
| Kaempferol | n.s. | n.s. | n.s. |
| Quercetin | * | * | n.s. |
| Genistein | ** | ** | n.s. |
| Caffeic Acid | *** | *** | n.s. |
| Daidzein | n.s. | n.s. | * |
| Estradiol | n.s. | n.s. | * |
| Cresol | n.s. | n.s. | n.s. |
| Nitrocatechol | *** | *** | ** |
| Phenolphthalein | * | * | n.s. |

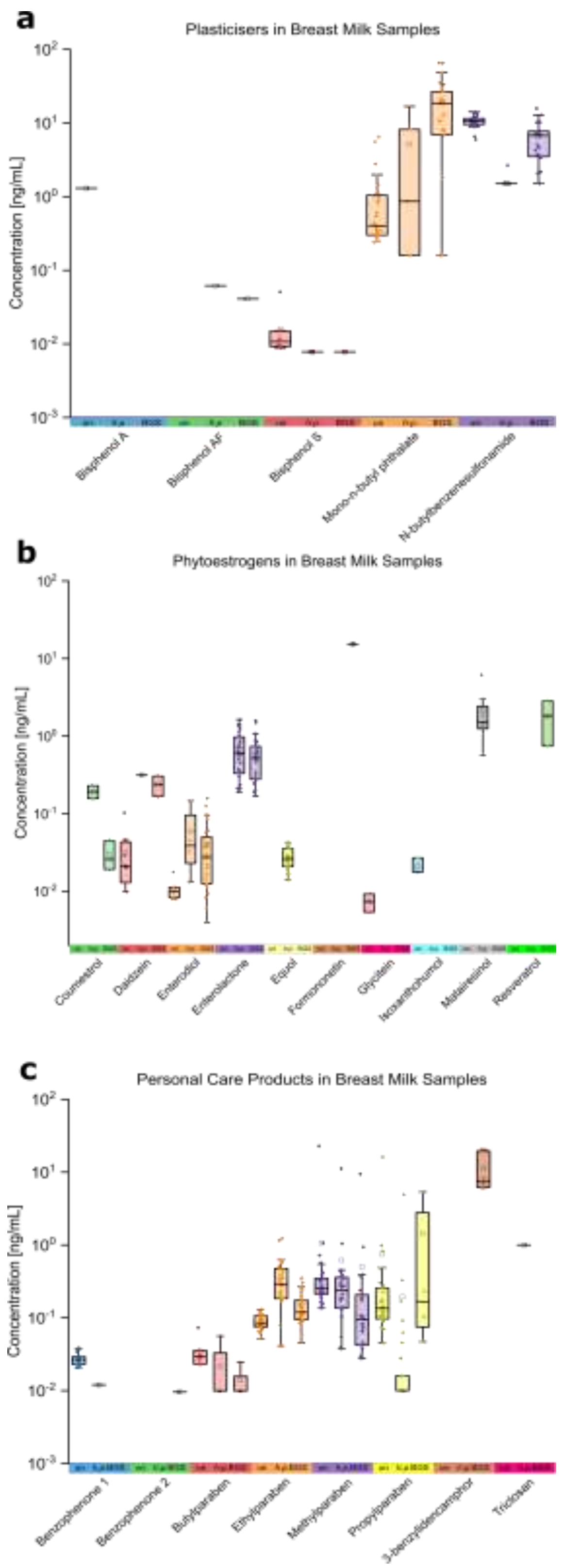

**Figure S 1.** Results of a subset of the long-term longitudinal breast milk study (n=30).
One mother collected her breast milk across 89 days postpartum for (a) plasticisers, (b) phytoestrogens, and (c) personal care products. These samples were measured without enzymatic hydrolysis (un), and with an enzymatic treatment of H. pomatia (h.p.) and BGS recombinant (BGS). The black line in the box represents the median line, the small square the mean, the box the 25–75 percentiles, and the whiskers the interquartile range. In the table, the minimum (min) and maximum (max) concentration in ng/mL are given together with the average (avg) and number of positive samples (n). The concentrations are displayed in a decadic logarithmic scale.
